# Effects of population- and seed bank noise on neutral evolution and efficacy of natural selection

**DOI:** 10.1101/168682

**Authors:** Lukas Heinrich, Johannes Müller, Aurélien Tellier, Daniel Živković

## Abstract

Population genetics models typically consider a fixed population size and a unique selection coefficient. However, population dynamics inherently generate noise in numbers of individuals and selection acts on various components of the individuals’ fitness. In plant species with seed banks, the size of both the above- and below-ground compartments present noise depending on seed production and the state of the seed bank. We investigate if this noise has consequences on 1) the rate of genetic drift, and 2) the efficacy of selection. We consider four variants of two-allele Moran-type models defined by combinations of presence and absence of noise in above-ground and seed bank compartments. Time scale analysis and dimension reduction methods allow us to reduce the corresponding Fokker-Planck equation to a one-dimensional diffusive Moran model. We first show that if the above-ground noise classically affects the rate of genetic drift, below-ground noise reduces the diversity storage effect of the seed bank. Second, we consider that selection can act on four different components of the plant fitness: plant or seed death rate, seed production or seed germination. Our striking result is that the efficacy of selection for seed death rate or germination rate is reduced by seed bank noise, whereas selection occurring on plant death rate or seed production is not affected. We derive the expected site-frequency spectrum reflecting this heterogeneity in selection efficacy between genes underpinning different plant fitness components. Our results highlight the importance to consider the effect of ecological noise to predict the impact of seed banks on neutral and selective evolution.

## 1. Introduction

Genetic drift and natural selection are prominent forces shaping the amount of genetic diversity in populations. In diploid dioecious organisms, natural selection can be decomposed in different components: 1) viability selection as the differential survival of the genotypes from zygotes to adults, 2) fecundity (or fertility) selection as the differential zygote production, 3) sexual selection as the differential success of the genotypes at mating, and 4) gametic selection as the distorted segregation in heterozygotes (Bundgaard and Christiansen, 1972; Clegg et al., 1978). In effect, population genetic models with discrete generations or with Malthusian parameter ignoring age-structure lump these components into one unique selection parameter. Experimental or genomic population studies describing thus changes in allele frequencies often fail to describe and dissect the respective effects of these selective modes. Several theoretical studies on fertility (Bodmer, 1965) or on sexual selection (Karlin and Scudo, 1969) as well as experimental work on animal (Prout, 1971b,a; Christiansen and Frydenberg, 1973) and plant (Clegg et al., 1978) populations have attempted to disentangle the respective influence of these selection components.

In age-structured populations, however, genetic drift and selection can act differently than predicted by discrete models (overview in the book by Charlesworth, 1994). The first type of age-structured models are simply obtained by individuals’ life span and reproduction overlapping several generations. Here, genetic drift acts equally on all individuals of all age classes at any generation. The magnitude of genetic drift is defined by the population size which can be fixed, or fluctuating following a logistic dynamic size constrained by the population carrying capacity. As an extreme type of overlapping generation model, the Moran model exhibits a rate of random genetic drift all but the same as the Wright-Fisher (WF) model up to a rescaling of the population size (see, e.g., the book by Ewens, 2004). Meanwhile, in age-structured populations, selection for fecundity and viability can show different outcomes, such as time to allele fixation and/or maintenance of alleles, compared to discrete models. This occurs if fecundity or longevity act at different ages of the structured population and under a logistic population size dynamic (Anderson and King, 1970; King and Anderson, 1971; Charlesworth and Giesel, 1972a,b). An interesting question arising from the current increasing availability of genomic data is whether in age-structured populations selection for fecundity can be disentangled from that of viability using population genomics statistics (such as the site-frequency spectrum, SFS).

A second type of age-structured model is obtained when considering that individuals may remain as dormant/quiescent structures spanning several generations. Quiescence in reproductive structure is in fact wide-spread, such that seeds or eggs can be persistent states that allow to buffer a variable environment (Evans and Dennehy, 2005). The time at which offspring germinates or hedges can be variable, such that only some of the offspring live in detrimental periods, and most likely at least some in a beneficial environment. Bacterial spores or lysogenic states of temperate phages are also examples of such a bet-hedging strategy. Seedbanks represent thus a storage of genetic diversity decreasing the probability of population extinction (Brown and Kodric-Brown, 1977), diminishing the effect of genetic drift (Nunney, 2002), slowing down the action of natural selection (Templeton and Levin, 1979; Koopmann et al., 2017) and favouring balancing selection (Tellier and Brown, 2009). The strength of the seed bank effect clearly depends on the organism under consideration. Plants and eggs have a somehow short live span compared to the average coalescent time, and we call these seed banks “weak”. Dormant states of bacteria, however, can last a longer time (many generations) even than the average coalescent time. These seed banks are called “strong”, and modelled in a similar way as weak selection: the time scale of the quiescent state is scaled by the inverse of the population size. In particular Blath et al. (2015, 2016) investigated in a series of papers strong seed banks, and find mathematically appealing results as deviations from the Kingman coalescent.

In the present paper, we focus on weak seed banks aiming at applications to plant or invertebrate species. In a seminal paper Kaj et al. (2001) investigated the effect of weak seed banks on the coalescent. The key parameter is the average time seeds spend in the bank, denoted here as 1 + *G*. Kaj et al. (2001) have obtained the Kingman *n*-coalescent rescaled by (1 + *G*)^−2^. It was subsequently shown that 1 + *G* can be estimated using polymorphism data and information on the census size of populations (Tellier et al., 2011). Furthermore, neglecting seed banks may yield distorted results for the inference of past demography using for example the SFS (Živković and Tellier, 2012). Interestingly, the effect of (weak) selection is enhanced by the slow-down of the time scale due to seed banks (Blath et al., 2013, 2016; Koopmann et al., 2017). The effect of genetic drift and weak natural selection on allele frequencies can be computed in a diffusion framework in a Moran model with deterministic seed bank (Koopmann et al., 2017). While the diffusion term, defining genetic drift is scaled by (1 + *G*)^−2^, matching the backward coalescent result of Kaj et al. (2001), the coefficient of natural selection in the drift term is multiplied only by (1 + *G*)^−1^. In biological terms, this means that the strength of selection, as defined by a unique selective coefficient, is enhanced by the seed bank compared to the effect of genetic drift, even though the time to reach fixation for an allele is increased (Koopmann et al., 2017).

We investigate two additions to this current body of theoretical literature on weak seed banks. First, we compute the effect of realistic models relaxing the hypotheses of a fixed size for the population above ground, and of deterministically large for the seed bank compartment. By doing so, we generate noise in the population size above or below ground. This extends the classic Moran or Wright-Fisher models, as the importance of the noise effect on population dynamics on the rate of genetic drift or selection is being recognized (Huang et al., 2015). However, this approach is faced with an additional difficulty: if two alleles are present in a population of a constant size, it is sufficient to keep track of the number of individuals for one allele only. In the case of a fluctuating population size with logistic dynamics constrained by the carrying capacity the state space becomes essentially two-dimensional. This more general case can still be examined via is a time scale analysis and a dimension reduction by singular perturbation approaches. These methods are applicable in the case where the population size becomes large (Parsons and Quince, 2007; Parsons et al., 2008; Kogan et al., 2014). Though in those papers (as in the present one) the arguments are used in a formal way, the validity of this approach is proven (Kuehn, 2015). This latter analysis reveals that the dynamics are well described by appropriate scaled diffusive Moran models. Second, we dissect the fitness of plants into four components, which can be possibly affected by genetic drift occurring above ground and in the seed bank. We compute classic population genetics results for neutral and selected alleles and derive the expected SFS for the alleles under different fitness components. Using our model of a weak seed bank (with or without additional noise), we find approximative, appropriate scaled diffusive Moran models. These rescaled Moran models reveal the effect of noise in the above-ground population (plants) as well as in the below-ground population (seeds). In particular, we find that genetic drift and selection are differently influenced by above and below-ground noise. Additionally, the selection coefficient of alleles involved in seed death rate or germination rate are reduced by seed bank noise, but not by above-ground noise, while the two others are not affected (plant death rate and seed production). Below-ground noise also reduces the seed bank storage effect of neutral genetic diversity.

## 2. Methods

The aim of the present study is to investigate the effect of above-ground/below-ground population noise on evolution in presence of seed banks. Noise in population size can be located either above ground (plant population), or below ground (seed bank or seed population). Traditionally, the plant above-ground population is assumed to consist of *N* individuals, *N* being fixed. A noisy substitute of this assumption is a logistic (fluctuating) population model (Anderson and King, 1970; King and Anderson, 1971; Charlesworth and Giesel, 1972a,b). For seeds, we find in the literature either the assumption of a finite, fixed number (Kaj et al., 2001) of seeds, or, a deterministic infinite seed density (Koopmann et al., 2017). In the latter, the assumption is that the number of seeds per plant is large enough, such that intrinsic noise is negligible. We suggest, here, an alternative model, in which each plant produces single seeds at time points that are distributed according to a Poisson process. The seeds also die (or loose their ability to germinate) after a random time. Thus this “fluctuating seed bank” assumption assumes stochastically varying seed bank size. We therefore consider four models defined by all possible combinations of fixed or logistic above-ground population and deterministic or fluctuating seed bank.

**Notation:** *X*_1,*t*_ denotes the number of allele-*A* plants. In a model with fixed above-ground population size *N*, the number of allele-*B* plants reads *X*_2,*t*_ = *N* – *X*_1,*t*_; in the logistic fluctuating version, we set up a stochastic process for *X*_1,*t*_ and *X*_2,*t*_. In that case, *N* does denote the carrying capacity of the population (*i.e.*, the maximal possible size). The average population size (conditioned on non-extinction) is *κN* for some *κ* ∈ (0, 1). *Y_t_* (*Z_t_*) always refers to the amount of allele *A* (allele *B*) seeds in the bank. For the deterministic seed bank, *Y_t_* (*Z_t_*) are real numbers that follow (conditioned on *X*_1,*t*_, *X*_2,*t*_) an ordinary differential equation (ODE). In the case of stochastically fluctuating seed banks, *Y_t_* (*Z_t_*) are non-negative integers, that follow a stochastic birth-death process.

We allow for weak natural selection, so that allele *B* has a disadvantage. The rates for allele *B* slightly differ from those for allele *A* individuals on a scale of 1/*N*, as it is usual for weak effects. The parameters of our models are summarized in Table 1.

**Table 1:** Parameters of the models.

| meaning | symbol (Allele $A$) | symbol (Allele $B$) |
| --- | --- | --- |
| death rate of plants | $\zeta$ | $\zeta (1 + \sigma_1/N)$ |
| death rate of seeds | $\mu$ | $\mu (1 + \sigma_2/N)$ |
| production rate of seeds | $\beta$ | $\beta (1 - \sigma_3/N)$ |
| germination rate of seeds (log. pop. only) | $\gamma$ | $\gamma (1 - \sigma_4/N)$ |

### 2.1. Fixed population size and deterministic seed bank

This model has been developed and discussed before (Koopmann et al., 2017). The total above-ground population has size *N*, and the transitions for the allele-*A* plant population *X*_1,*t*_ ∈ {0, …, *N*} given the seed densities *Y_t_* and *Z_t_* ∈ ℝ_+_ are presented in Table 2.

**Table 2:** Possible transitions and their rates.

| event | offset | rate |
| --- | --- | --- |
| death of $A$, birth of $B$ | $X_{1,t} \mapsto X_{1,t} - 1$ | $\zeta X_{1,t} Z_t / (Y_t + Z_t)$ |
| death of $B$, birth of $A$ | $X_{1,t} \mapsto X_{1,t} + 1$ | $\zeta (1 + \sigma_1/N) (N - X_{1,t}) Y_t / (Y_t + Z_t)$ |

The dynamics of seeds, in turn, is given by a Davis’ piecewise deterministic process (Davis, 1984),

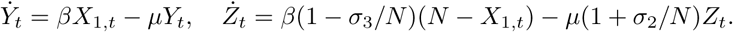

### 2.2. Logistic population dynamics and deterministic seed bank

For the logistic model, we do not couple death and birth of a plant as it is usually done in Moran-type models to keep the total population size constant. We generalize the logistic dynamics as investigated, e.g., by Nasell (2011), or Parsons and Quince (2007); Parsons et al. (2008) for the situation at hand and separate birth and death events. If the seed densities *Y_t_*, *Z_t_* ∈ ℝ_+_ are given, the transitions for *X*_1,*t*_, *X*_2,*t*_ ∈ {0, …, *N*}, *X*_1,*t*_ + *X*_2,*t*_ ≤ *N*, read as summarized in Table 3.

**Table 3:**
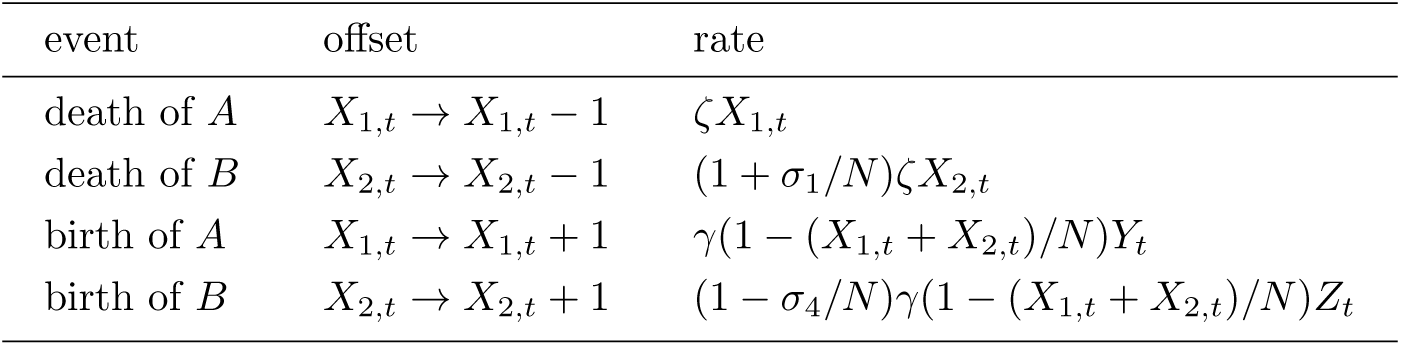
Possible transitions and their rates.

| event | offset | rate |
| --- | --- | --- |
| death of $A$ | $X_{1,t} \rightarrow X_{1,t} - 1$ | $\zeta X_{1,t}$ |
| death of $B$ | $X_{2,t} \rightarrow X_{2,t} - 1$ | $(1 + \sigma_1/N)\zeta X_{2,t}$ |
| birth of $A$ | $X_{1,t} \rightarrow X_{1,t} + 1$ | $\gamma(1 - (X_{1,t} + X_{2,t})/N)Y_t$ |
| birth of $B$ | $X_{2,t} \rightarrow X_{2,t} + 1$ | $(1 - \sigma_4/N)\gamma(1 - (X_{1,t} + X_{2,t})/N)Z_t$ |

Conditioned on *X*_1,*t*_ and *X*_2,*t*_, the dynamics of seeds are again deterministic,

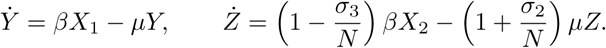

### 2.3. Fixed population size and fluctuating seed bank

For the above-ground population we return to a fixed population size, s.t. it is sufficient to follow *X*_1,*t*_ as *X*_2,*t*_ = *N* – *X*_1,*t*_. In the present model we address the noise in the number of seeds, *Y_t_*, *Z_t_* ∈ ℕ_0_. The seeds follow a stochastic birth-death process, where the death rate is kept constant, and the birth rate is proportional to the number of corresponding above-ground plants. We obtain the transitions summarized in Table 4.

**Table 4:** Possible transitions and their rates.

| event | offset | rate |
| --- | --- | --- |
| death of $A$, birth of $A$ | $(X_{1,t}, Y_t) \rightarrow (X_{1,t}, Y_t - 1)$ | $\zeta X_{1,t} Y_t / (Y_t + Z_t)$ |
| death of $A$, birth of $B$ | $(X_{1,t}, Z_t) \rightarrow (X_{1,t} - 1, Z_t - 1)$ | $\zeta X_{1,t} Z_t / (Y_t + Z_t)$ |
| death of $B$, birth of $A$ | $(X_{1,t}, Y_t) \rightarrow (X_{1,t} + 1, Y_t - 1)$ | $(1 + \sigma_1/N)\zeta(N - X_{1,t})Y_t / (Y_t + Z_t)$ |
| death of $B$, birth of $B$ | $(X_{1,t}, Z_t) \rightarrow (X_{1,t}, Z_t - 1)$ | $(1 + \sigma_1/N)\zeta(N - X_{1,t})Z_t / (Y_t + Z_t)$ |
| birth of $A$-seed | $Y_t \rightarrow Y_t + 1$ | $\beta X_{1,t}$ |
| death of $A$-seed | $Y_t \rightarrow Y_t - 1$ | $\mu Y_t$ |
| birth of $B$-seed | $Z_t \rightarrow Z_t + 1$ | $(1 - \sigma_3/N)\beta(N - X_{1,t})$ |
| death of $B$-seed | $Z_t \rightarrow Z_t - 1$ | $(1 + \sigma_2/N)\mu Z_t$ |

### 2.4. Logistic population dynamics and fluctuating seed bank

The last model incorporates logistic growth and a stochastically fluctuating seed bank. This model is an obvious combination of the last two models,

**Table 5:** Possible transitions and their rates.

| event | offset | rate |
| --- | --- | --- |
| death of $A$ | $X_{1,t} \rightarrow X_{1,t} - 1$ | $\zeta X_{1,t}$ |
| death of $B$ | $X_{2,t} \rightarrow X_{2,t} - 1$ | $(1 + \sigma_1/N)\zeta X_{2,t}$ |
| birth of $A$ | $(X_{1,t}, Y_t) \rightarrow (X_{1,t} + 1, Y_t - 1)$ | $\gamma(1 - (X_{1,t} + X_{2,t})/N)Y_t$ |
| birth of $B$ | $(X_{2,t}, Z_t) \rightarrow (X_{2,t} + 1, Z_t - 1)$ | $(1 - \sigma_4/N)\gamma(1 - (X_{1,t} + X_{2,t})/N)Z_t$ |
| birth of $A$-seed | $Y_t \rightarrow Y_t + 1$ | $\beta X_{1,t}$ |
| death of $A$-seed | $Y_t \rightarrow Y_t - 1$ | $\mu Y_t$ |
| birth of $B$-seed | $Z_t \rightarrow Z_t + 1$ | $(1 - \sigma_3/N)\beta X_{2,t}$ |
| death of $B$-seed | $Z_t \rightarrow Z_t - 1$ | $(1 + \sigma_2/N)\mu Z_t$ |

### 2.5. Strategy for the analysis of the models

The aim of the analysis is the reduction of the four models to a one-dimensional diffusive Moran model representing the fraction of allele-*A* individuals within the population. As the details of the analysis are tedious, we present them in detail in the appendix and only outline the basic idea in the present section.

The strategy of the analysis differs for the first model (fixed population size, deterministic seed bank) and the other three models. The reason is that there is one single stochastic state variable *X*_1,*t*_ in the first model, so that if we know the history of *X*_1,*t*_, the state of the seed bank is known. No dimension reduction method is thus required. We basically can use the approach of Koopmann et al. (2017) to derive the diffusive Moran model. However, we outline in the appendix an argument based on a small-delay approximation, as this route seems to provide an appealing short-cut: if ecological time *t* is not considered but rather the evolutionary time *τ* = *t*/*N* (*N* large), the delay of a weak seed bank is small. In this case, the solution can be expanded w.r.t. the delay. As a result, the seed bank can be removed from the stochastic process and replaced by appropriately rescaled parameters.

The three other models have two, three or four stochastic state variables. Methods of dimension reduction are required to obtain a one-dimensional diffusive Moran model. The key insight here is that any realization saddles fast on a one-dimensional manifold. If we consider, e.g., the deterministic version of the logistic population dynamics with stochastically fluctuating seed bank, we find, according to arguments by, e.g., Ethier and Kurtz (1986), for the deterministic limit as *N* → ∞ (with *x_i_*(*t*) = *X*_*t*,*i*_/*N*, *y*(*t*) = *Y_t_*/*N*, *z*(*t*) = *Z_t_*/*N*)

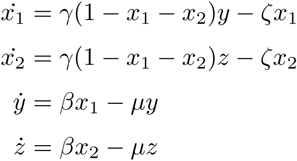

It is straightforward to show that a line of stable equilibria, the so-called coexistence line, exists:

#### Proposition 2.1.

*Let ϑ*:= (*β* – *ζ*)/*μ* > 0, *κ*:= (*γϑ* – *ζ*)/(*γϑ*) ∈ [0, 1]. *Then*, *there is a line of stationary points in* [0, *κ*]^2^ × 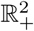 *given by*

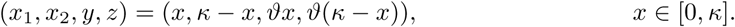

*The line of stationary points is transversally stable (locally and globally).*

It turns out that the stochastic process rapidly approaches this line of equilibria, and performs a random walk close to it (see figure 1). The analysis reveals that the distribution on a transversal cut is just a normal distribution with a variance of 𝒪(1/*N*). Along the line of stationary points, however, the realizations will move according to a one-dimensional Moran process. In order to reveal this structure, we first use a large population limit to obtain a Fokker-Planck/Kolmogorov forward equation for the full process. Then, we use singular perturbation methods as described, e.g., in Kogan et al. (2014) or Kuehn (2015) to perform the dimension reduction to the one-dimensional Moran model.

**Figure 1:**
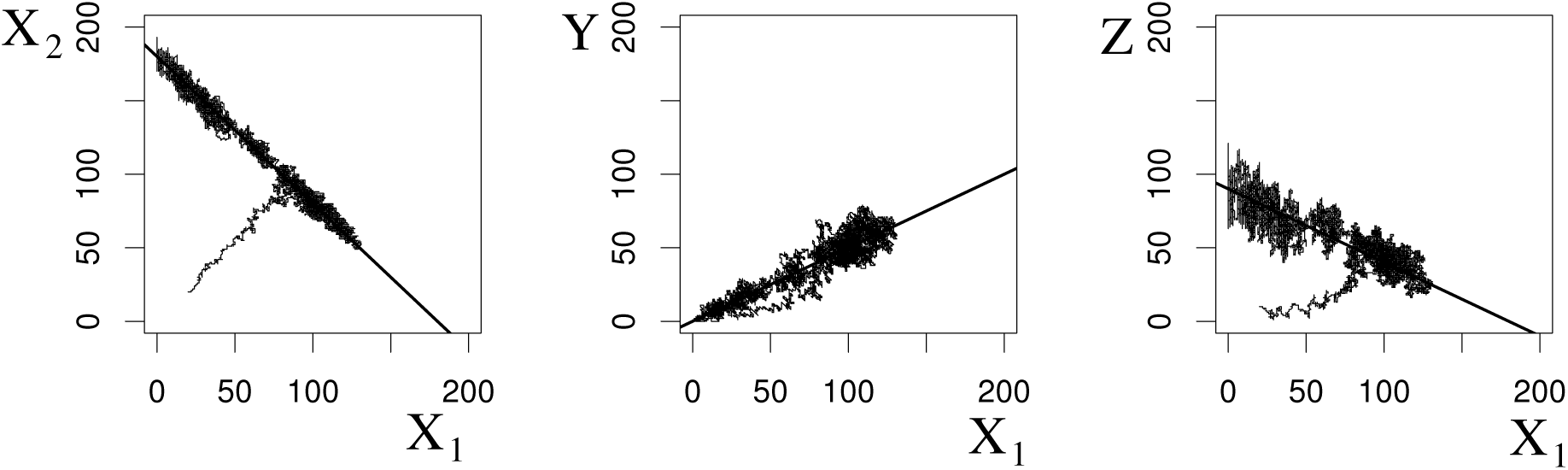
Simulated trajectory for model 2.3, (*left*) *A*-plants vs *B*-plants, (*center*) *A*-plants vs *A*-seeds, (*right*) *A*-seeds vs *B*-seeds, *N* = 200, *θ* = 0.5, *κ* = 0.9. The solid line indicates the coexistence line.

## 3. Results

### 3.1. Timescales for different seed bank models

For all of our models, the resulting one-dimensional Fokker-Planck equation assumes the form of a diffusive Moran-model with weak selection,

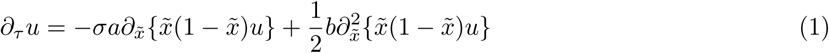

where *a* and *b* describe the speed of selection and genetic drift, respectively. The term *σ* represents selective coefficients and is a generic parameter for notation including *σ_i_*, *i* = 1, …, 4. In order to formulate the results for *a* and *b*, let us introduce three composite parameters: *G* = *ζ*/*μ* is the number of plant generations a seed survives on average; *Y* = *β*/*ζ* is the average number of seeds produced by a plant; *κ* already defined in Proposition 2.1 is the average above-ground population size of the logistic model. Furthermore, for deterministic and fluctuating seed banks, we respectively denote (1 + *G*)^−1^ and (1 + (1 – 1/*Y*)*G*)^−1^ as 𝒢, which can be seen as the number of plant generations that seeds survive on average corrected by the size of the seed bank. Using these abbreviations, the parameters *a*, *b* and *σ* for our four models are summarized in Table 6.

**Table 6:** Drift, diffusion, and selection coefficients for the different population/seed bank models.

| model/scale term | $a(\text{selection})$ | $b(\text{genetic drift})$ | $\sigma$ (selection coeff.) |
| --- | --- | --- | --- |
| fix. pop., det. s.b. | $\zeta \mathcal{G}^{-1}$ | $2\zeta \mathcal{G}^{-2}$ | $\sigma_1 + \sigma_2 + \sigma_3$ |
| fix. pop., fluct. s.b. | $\zeta \mathcal{G}^{-1}$ | $2\zeta \mathcal{G}^{-2}$ | $\sigma_1 + (1 - 1/Y)\sigma_2 + \sigma_3$ |
| log. pop., det. s.b. | $\zeta \mathcal{G}^{-1}$ | $2\zeta/\kappa \mathcal{G}^{-2}$ | $\sigma_1 + \sigma_2 + \sigma_3 + \sigma_4$ |
| log. pop., fluct. s.b. | $\zeta \mathcal{G}^{-1}$ | $2\zeta/\kappa \mathcal{G}^{-2}$ | $\sigma_1 + \sigma_2 + \sigma_3 + (1 - 1/Y)\sigma_4$ |

The basic seed bank model (fixed population size, deterministic seed bank) and the standard Moran model without seed bank can be used as reference models. A seed bank slows down the time scale of selection as well as that of genetic drift (Koopmann et al., 2017), where selection is less affected (by a factor of (1 + *G*)^−1^) than genetic drift (by a factor of (1 + *G*)^−2^). We find that fluctuations in the above-ground population and in the seeds have different effects.

Fluctuations in the seed number reduce the storage effect of seed banks. The additional noise yields a reduction of the effective time a seed spends within the seed bank, and thus increases the rate of genetic drift. For *Y* → ∞ (noise in seed bank tends to zero), we obtain the result for the deterministic seed bank, for *Y* → 1 (noise is maximized), the model converges towards the standard Moran model without seed bank. Note that *Y* is not the average number of seeds per plant directly measured but the effective number of seeds per plants. For example, a certain fraction of seeds might be getting lost due to other environmental reasons (abiotic or biotic factors) than their intrinsic mortality. These seeds do not contribute to the bank.

The noise in the above-ground population only affects genetic drift and does not appear in the selection term. This result reflects that the actual competition between alleles *A* and *B* only happens above ground. Nonlinear terms in the transition rates only appear in the birth term of the plants. By increasing solely genetic drift, the above-ground noise can counteract the amplification of selection by seed banks.

Our first result is somehow expected. All mutations are affected in the same way by the above-ground noise. Our second result is, however, unexpected. Mutations for some fitness components (mortality of seeds, *σ*_2_, and germination ability, *σ*_4_) show reduced selection while this is not the case for selective coefficients of other fitness components (*σ*_1_ and *σ*_3_). This means that if the number of seeds per plant is not too large, beneficial mutations in the mortality of seeds (*σ*_2_) have a reduced chance to reach fixation compared with a beneficial mutation for the production of seeds (*σ*_3_).

### 3.2. Site-frequency spectrum (SFS)

The SFS is a commonly used statistics for the analysis of genomewide distributed SNPs. It is defined as the distribution of the number of times a mutation is observed in a population or a sample of *n* sequences conditional on segregation. Mutations are assumed to follow the infinitely-many sites assumption (Kimura, 1969) at rate *θ* = 2*N ν* (for a haploid population as assumed herein), where *ν* is the mutation rate per generation at independent sites. Mutations are allowed to occur in plants and seeds, but the following results can be easily adapted to the scenario, where mutations may only arise in plants. Assuming that each mutant allele marginally follows the diffusion model specified in (1), the proportion of sites at equilibrium, where the mutant frequency is in (*y*, *y* + *dy*), can be obtained as (e.g., Koopmann et al. 2017)

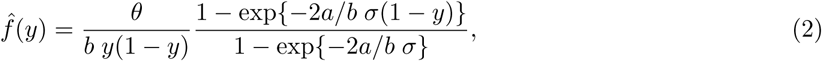

where *a*, *b* and *σ* ≠ 0 are given in Table 6. The sample SFS at equilibrium can be immediately obtained from (2) via binomial sampling as

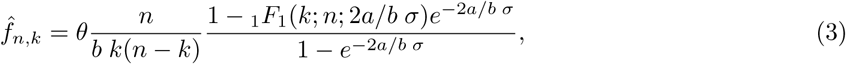

where _1_*F*_1_ denotes the confluent hypergeometric function of the first kind (Abramowitz and Stegun, 1964). The neutral versions of (2) and (3) are respectively given by *f̂*(*y*) = *θ*/(*by*) and *f̂*_*n*,*k*_ = *θ*/(*bk*).

Fig. 2 indicates the striking effects of the seed bank noise for the model with logistic population dynamics and fluctuating seed banks. When the noise is low (*Y* high) the number of segregating sites is expected to be high, and mutations involved in the selection coefficients *σ*_1_ and *σ*_4_ lead to similar SFS. The SFS show the typical U-shape expected under positive pervasive selection. If, however, *Y* becomes small, the number of segregating sites decreases, and the selection coefficient *σ*_1_ shows a U-shaped SFS while mutations under *σ*_4_ do not.

**Figure 2:**
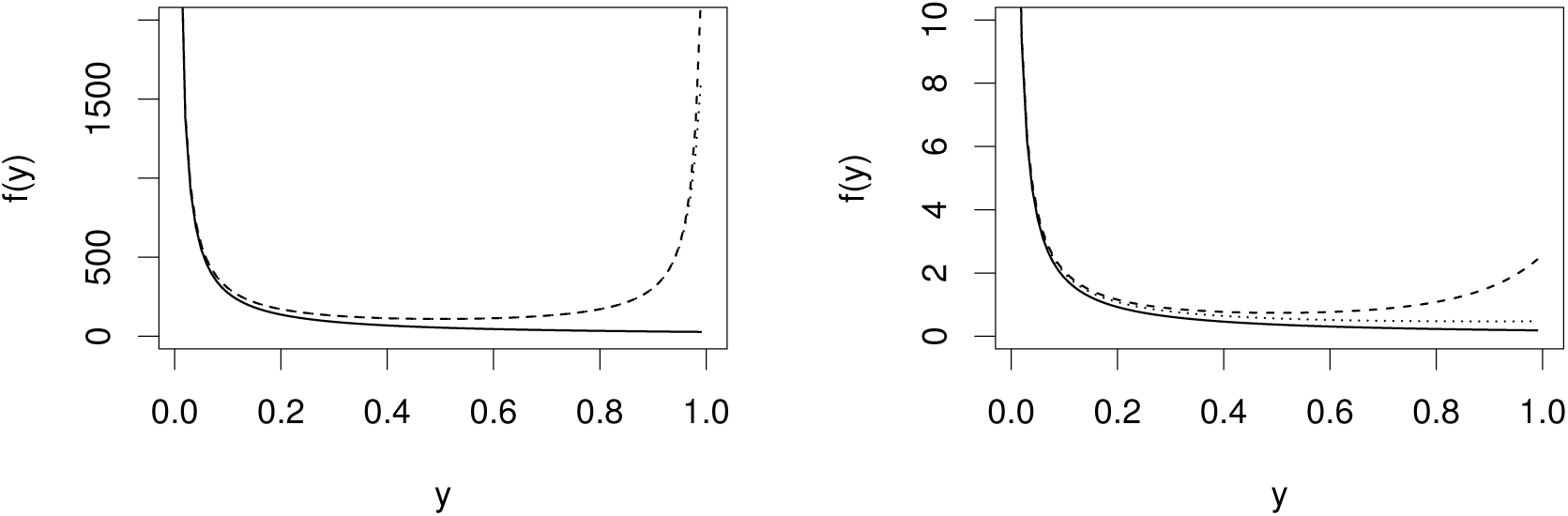
Continuous SFS, neutral (solid) vs. selection (dashed, *σ*_1_ = 50, *σ*_2_ = *σ*_3_ = *σ*_4_ = 0) resp. (dotted, *σ*_1_ = *σ*_2_ = *σ*_3_ = 0,
*σ*_4_ = 50). Further parameters: *μ* = 0.25, *ζ* = 1, *γ* = 1.5; (left) *β* = 1.2, *G* = 4, *Y* = 1.2, (right) *β* = 2.4, *G* = 4, *Y* = 2.4.

## 4. Discussion

We considered four models to investigate the effect of combined noise in the above-ground population and below-ground seed bank. In all four cases time scale arguments allow us a reduction to a diffusive Moran model. Our results extend the findings that a seed bank without noise yields a change of time scale in selection and genetic drift that amplifies the effect of weak selection (Koopmann et al., 2017). The first main result of this present work is that there is no direct interaction between the noise above ground and below ground. The above-ground noise increases the effect of genetic drift compared to a fixed above-ground population size, but does not affect selection. One can propose a new definition of the effective population size *N_e_* describing the change in genetic drift due to this noise. This allows us to redefine the evolutionary time scale. As a result of this procedure the parameter *κ*, which represents the reduction of the average total population size and indicates the increase of the above-ground noise by the logistic population dynamics, appears as a factor in front of the selection term in the Moran model. However, note that in this notation we also find that the terms for below-ground and above-ground noise are multiplied indicating their independence.

The second result is that the below-ground noise affects the scaling of time for both the selection and the genetic drift term. We introduce the concept of the mean effective number of seeds per plant in a similar definition to the effective population size (Wright, 1931). If this average seed number tends to infinity, we recover the effect of a deterministic seed bank (no noise), and if this number tends to one, the noise of the seed bank accelerates the time scale such that the seed bank has no effect at all. The magnitude of noise in the seed bank is thus tuned between the two extreme cases of “no seed bank” (minimal effect) and “deterministic seed bank” (maximal effect).

The third result of the present study is the insight that below-ground noise may affect the four plants’ fitness components of viability and fecundity differentially. If the below-ground noise is large, the selective effect of mutations involved in seed death and germination may even be cancelled out. Other mutations involved in fitness traits of the above-ground population, such as plant death or seed production are not affected. In other words, while finite size and noise above ground do not affect selection, noise and finite size of the seed bank do change the selection coefficients. In biological terms we interpret this result as follows. Above ground, the fate of an allele under positive selection is classically determined by the strength of genetic drift, which depends on the population size (the diffusion term in the Fokker-Planck equation) compared to the strength of selection which depends on the selection coefficient (the drift term in the forward model). Under a deterministic seed bank (as in the present models and in Koopmann et al. (2017)), selection is efficient because it occurs on plants, when they are above ground, with a probability (1 + *G*)^−1^ and genetic drift occurs on a coalescent scale of (1 + *G*)^−2^. Any change in the allele frequencies above ground translates directly into the deterministic seed bank, just with a small time delay. However, when the seed bank has a fluctuating finite size, the strength of selection on seed fitness (the coefficients *σ*_2_ and *σ*_4_) is decreased by the noise in the seed compartment. This occurs because selection and genetic drift in the bank occur at every generation and not only when seeds germinate.

This observation implies that competition experiments, which measure and compare the effect of mutations and/or allow to determine selection coefficients on the ecological time scale can hardly be used to extrapolate to the evolutionary time scale. Indeed, even if two mutations seem to have an equivalent value for the plant fitness in a competition experiment, the presence of seed bank noise may lead to different evolutionary outcomes.

We finally discuss two ways of using genome polymorphism data to estimate seed bank parameters as well as the selection coefficients. We can attempt to infer the seed bank parameters based on neutral genetic diversity under the idealized conditions that the sample SFS reached an equilibrium as *f̂*_*n*,*k*_ (*i.e.*, there is no recent demographic impact) and that we can measure or estimate the population mutation rate *θ* and the death rate of plants *ζ*. It turns out that 𝒢 (the number of plant generations a seed survives on average) and *κ* (the average above-ground population size) can then be identified, so that it is possible to disentangle the effects of the seed bank from those of the above-ground noise on neutral evolution by utilizing *f̂*_*n*,*k*_. However, we cannot identify the below-ground noise, as *G* and *Y* only appear in the composite parameter 𝒢. Note that if we only have information about the relative sample SFS *f̂*_**n*,*k**_/∑_*i*_ *f̂*_*n*,*i*_, the multiplicative constant in *f_n_*,· cancels out, and therefore only *κ* 𝒢 and the combined effect of above- and below-ground noise can be estimated. Extending the work by Tellier et al. (2011), we suggest that the number of plant generations a seed survives on average can be estimated from the absolute SFS using for example a Bayesian inference method with priors on the census size of the above-ground population and the death rate of plants.

We may also aim to infer the selection coefficients underpinning the various plants’ fitness components, namely the fecundity and viability of plants or seeds. Our expected SFS show that the behavior of the seed fitness coefficients can be affected by the below-ground noise. So, we predict that populations with a small sized seed bank should exhibit less selection signatures on genes related to seed fitness compared to populations with a larger seed bank compartment. With the abundance of gene expression, molecular and Gene Ontology data, it becomes feasible to group genes by categories of function or pathways, for example involved in seed germination or seed integrity, viability and seed dormancy (e.g., Righetti et al., 2015). These groups of genes would correspond to the different plants’ fitness components investigated in this study. Our prediction is thus that the SFS and the inferred distribution of selective effects (e.g., using the method by Eyre-Walker and Keightley, 2009) would reflect the differential selection on these fitness components. Our results differ thus from those of classic age-structured populations by the overlap of generations, as the seed bank can present its own rate of genetic drift. Moreover, selection acts differently above ground and below ground on the different plants’ fitness components, which may allow us to disentangle their effect on the overall selection coefficient.

## Acknowledgements

*This research is supported in part by Deutsche Forschungsgemeinschaft grants TE 809/1 (AT) and STE 325/14 from the Priority Program 1590 (DZ).*

## Appendix Analysis of the Models

We present the conceptions for the analysis of the model of a fixed population size and a deterministic seed bank as well as for logistic population dynamics and a deterministic seed bank in detail. Note that the computations for the remaining two models are similar but even more extensive, so that we do not present them in full length but only mention the results of the main steps. The computations for the dimension reduction have been checked using the computer algebra package MAXIMA (Maxima, 2014) (see the available supplementary files).

## A.1. Fixed population size and deterministic seed bank

To keep the demonstration short, we present a nice but heuristic argument using the idea of a short delay approximation (Guillouzic et al., 1999). A more in-depth analysis of the model for the cases *σ*_2_ = *σ*_3_ = 0 can be found in Koopmann et al. (2017). Using the variation-of constant formula, we find

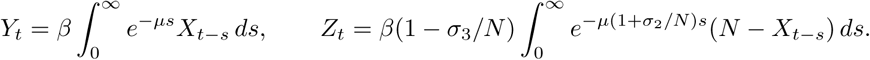

Let *x_t_* = *X*_1,*t*_/*N* and *ε*^2^ = 1/*N*. Then,

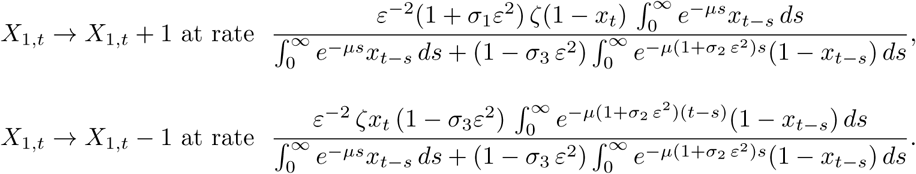

With standard arguments, we obtain a stochastic delay differential equation (SDDE) at the evolutionary time scale *τ* = *t ε*^2^ as

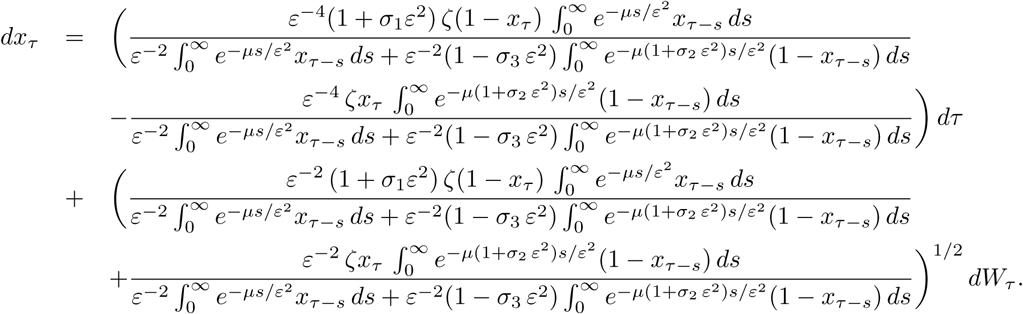

We aim a small delay approximation. Therefore, we note that for a function Φ(*t*), which is sufficiently smooth and bounded, we have (for *μ* > 0)

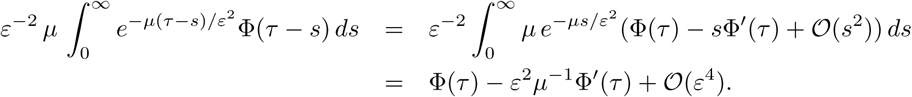

Thus, at a formal level,

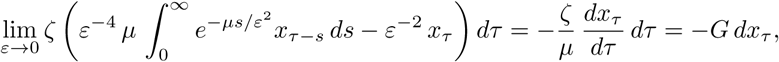

where *G* = *ζ*/*μ*. Note that this equation has to be interpreted in terms of the Euler-Maruyama approximation of an SDDE, where differential quotients are replaced by difference quotients. If we add

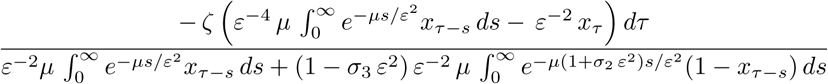

to both sides of the SDDE and let *ε* → 0, we obtain

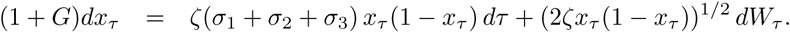

This equation yields the desired result with *σ* = *σ*_1_ + *σ*_2_ + *σ*_3_.

## A.2. Logistic population dynamics and deterministic seed bank

Let *p*_*i*,*j*_ (*k*, *l*, *t*) = ℙ(*X*_1,*t*_ = *i*, *X*_2,*t*_ = *j*, *Y_t_* ∈ (*k*, *k* + *dk*), *Z_t_* ∈ (*l*, *l* + *dl*)) be the joint probability density of the resulting stochastic process (with discrete *i*, *j* and continuous *k*, *l*). The corresponding master equation reads:

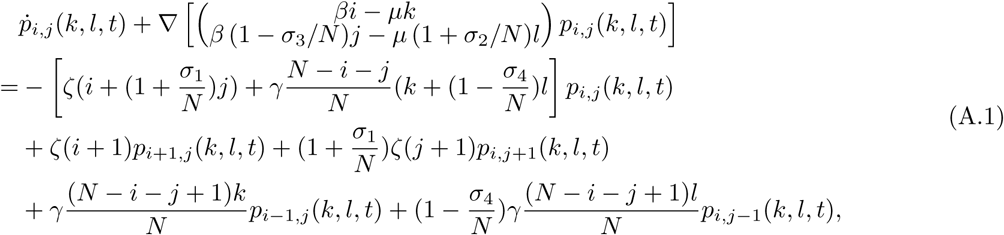

where the operator ∇ acts with respect to continuous state space variables *k*, *l*. Standard arguments yield the Fokker-Plank-approximation for large populations, where *x*_1_ = *i*/*N*, *x*_2_ = *j*/*N*, *y* = *k*/*N* and *z* = *l*/*N*, as

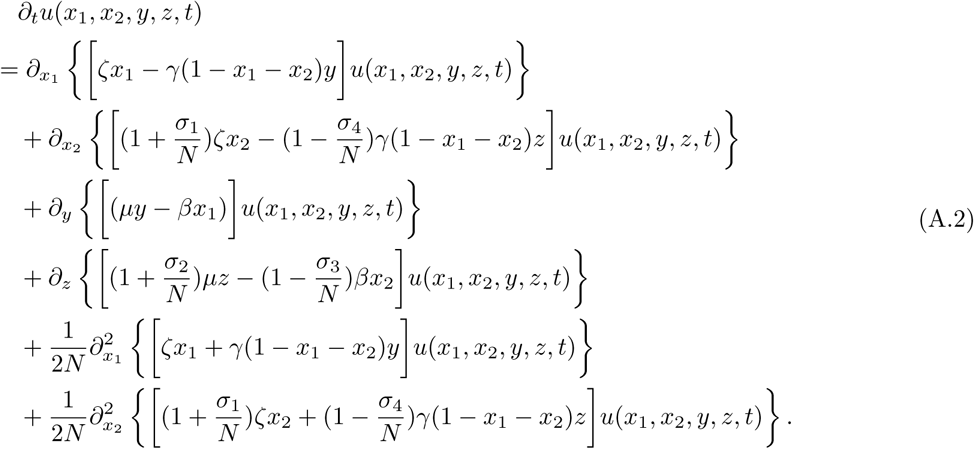

Note that the second order noise terms are solely due to *x*_1_ and *x*_2_; no noise is added by the seed bank variables *y* and *z*.

## A.2.1. Deterministic model

The corresponding deterministic model (drift terms only) yields the ODEs

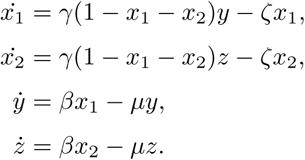

The lifetime reproductive success (or basic reproduction number) of a plant reads *R*_0_ = *β γ*/(*μζ*). If *R*_0_ > 1, the plant population can persist. Under this condition, there is a line of stationary solutions:

### Proposition A.1.

*Assume R*_0_ > 0. *Let ϑ*:= *β*/*μ*, *κ*:= (*γϑ* – *ζ*)/*γθ*. *Then*, *κ* ∈ [0, 1], *and there is a line of stationary points in* [0, *κ*]^2^ × 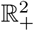 *given by*

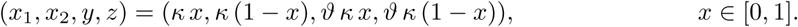

*The line of stationary points is transversally stable (locally and globally).*

### Proof

It is straightforward to check that the line indicated above consists indeed of stationary points with non-negative values. The local stability is a consequence of Hartman-Grobman and the analysis of the Jacobian (due to the block structure of this matrix the eigenvalues can be stated explicitly). Note that one eigenvalue necessarily is zero with an eigenvector pointing in the direction of the line of stationary points. Now, to show global stability, we prove that the system will approach the equilibrium line from any starting point. We first respectively denote *P* = *x*_1_ + *x*_2_ and *S* = *y* + *z* as the total plant and seed populations and consider the resulting reduced system

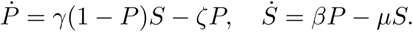

Here we use that we only consider weak selection: all selection effects tend to zero for *N* → ∞, and hence the ODE describes the neutral case. The divergence of this system is negative, and thus the combination of the theorems of Bendixon-Dulac and Pointcaré-Bendixon imply that trajectories (*P*, *S*) tend to stationary points. This observation yields the desired global stability.

Note that in equilibrium, *x*_1_ + *x*_2_ = *κ*. That is, *κ N* represents the average above-ground population size of the model conditioned on non-extinction.

## A.2.2. Dimension reduction by time scale analysis

The computations in this section follow closely the calculations in the paper of Kogan et al. (2014) to perform a dimension reduction by a time scale analysis. First, new local variables for the boundary layer around the equilibrium line are defined as

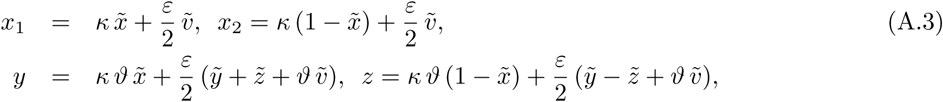

where

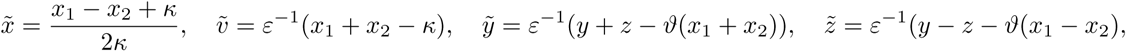

and *ε*^2^ = 1/*N*. For the transformed density *ρ*(*x̃*, *ṽ*, *ỹ*, *z̃*, *t*; *ε*) = *u*(*x*_1_, *x*_2_, *y*, *z*, *t*), we find

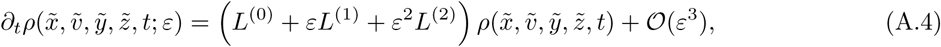

with linear differential operators

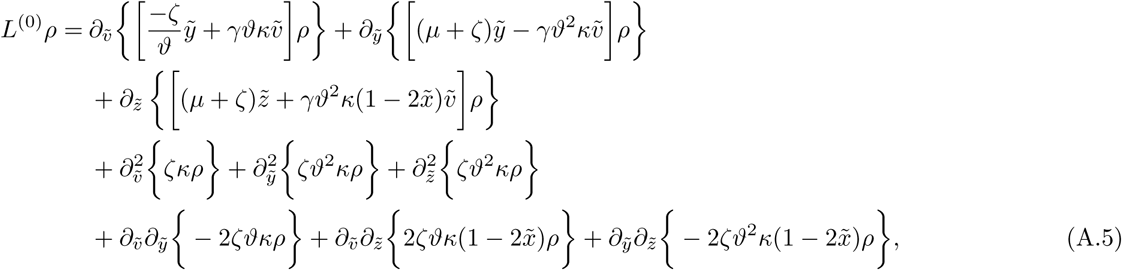

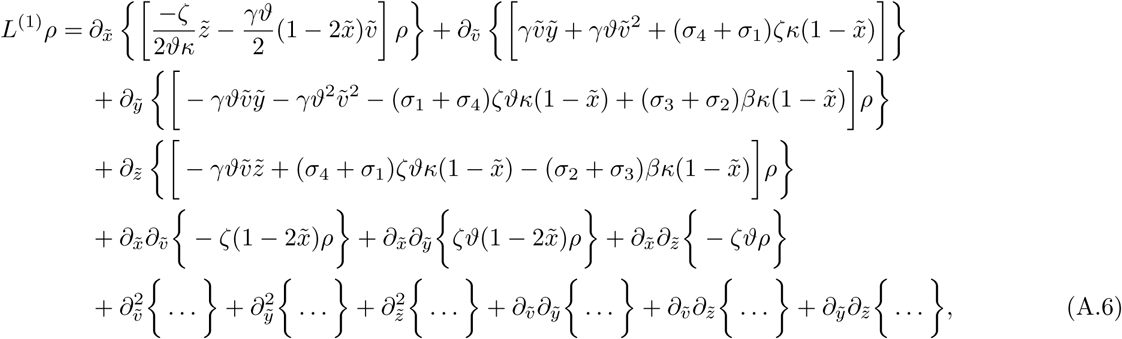

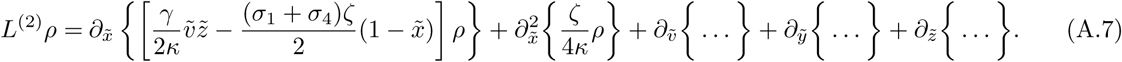

We employ a time scale separation and focus on a solution evolving on the slow time *τ* = *ε*^2^*t* = *t*/*N* using the Ansatz

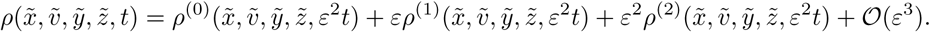

Plugging this into (A.4) and comparing same order terms, we have

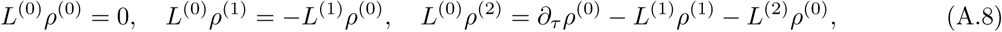

which indicates that *ρ*^(0)^ can be written as

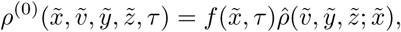

with a time independent normal distribution *ρ̂*(*ṽ*, *ỹ*, *z̃*; *x̃*) in *ṽ*, *ỹ*, *z̃* that satisfies *L*^(0)^*ρ̂* = 0. The function *f*(*x̃*, *τ*) modifies *ρ̂*(*ṽ*, *ỹ*, *z̃*; *x̃*) and represents the time evolution of *ρ*^(0)^ along the line of stationary points. Integrating the last equation of (A.8) from –∞ to ∞ with respect to *ṽ*, *ỹ*, *z̃* - the left hand side is a total derivative w.r.t. variables of integration and becomes zero - yields the evolution equation

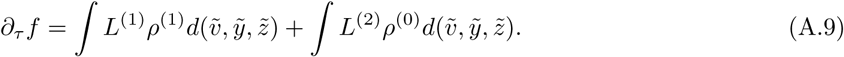

We start by computing the second integral. Since all terms that are full derivatives w.r.t. *ṽ*, *ỹ* or *z̃* vanish

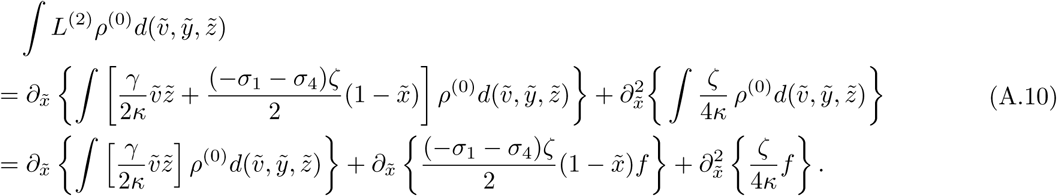

We take a closer look at the first integral. We aim to write *ṽz̃* = (*L*^(0)^)^+^*h*^+^ – *g*(*x̃*) for suitable *h*^+^, *g*, where (*L*^(0)^)^+^ is the adjoint of *L*^(0)^. If we can do so, then

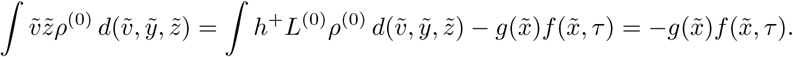

To identify *h*^+^ and *g*, we reduce the problem to linear algebra. Define the finite-dimensional vector space (for *k* ∈ ℕ)

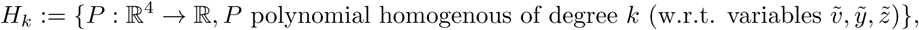

so that, e.g., *γ*/2*κ ṽz̃* ∈ *H*_2_, while *γ*/2*κ*/; *ṽz̃* + *c* ∉ *H*_2_ for *c* ∈ ℝ ∖ {0}. Examining (*L*^(0)^)^+^, we find

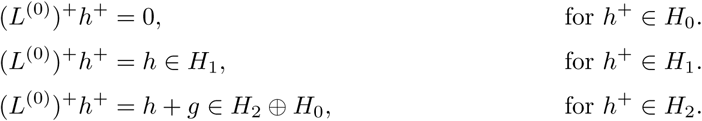

In particular the last observation (*L*^(0)^)^+^*H*_2_ → *H*_2_ ⊕ *H*_0_, *h*^+^ ↦ (*h*, *g*) allows to define an operator *M*: *H*_2_ → *H*_2_, *h*^+^ ↦ *h*. To simplify notation, we identify vectors w.r.t. a given basis in *H_k_* and the corresponding polynomials.

### Proposition A.2.

*W.r.t. the canonical basis* (*ṽ*^2^, *ỹ*^2^, *z̃*^2^, *ṽỹ*, *ṽz̃*, *ỹz̃*), *the operator M has the representation*

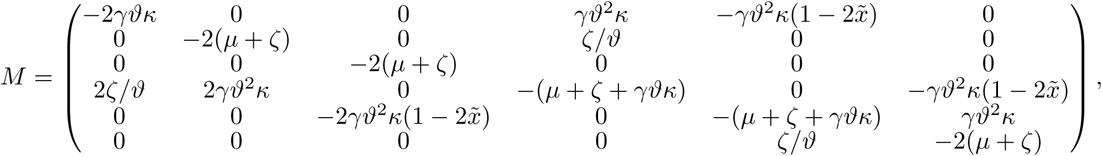

*and is invertible*, *if all parameters are positive. Let h* = (*a*_1_(*x̃*), …, *a*_6_(*x̃*))^*T*^ ∈ *H*_2_. *Then*, (*L*^(0)^)^+^(*h*) = *M h* + *g*(*x̃*), *where g*(*x̃*) ∈ *H*_0_ *is uniquely defined by g*(*x̃*) = *Ĝ*(*x̃*)*h and Ĝ*(*x̃*) *denoting the row*-*vector*

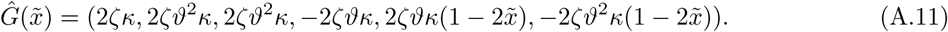

*For h̃* ∈ *H*_2_, *we find* (*L*^(0)^)^+^(*M*^−1^*h̃*) = *h̃* + *g*(*x̃*) *with g*(*x̃*) = *Ĝ*(*x̃*) *M*^−1^*h̃*, *so that*

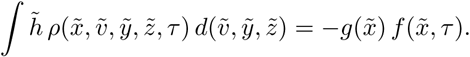

### Proof

It is straightforward to obtain the representation of *M* and *g* by applying *L*^(0)^ to the elements of the basis given above. Using, e.g., Gauß-elimination, we find that *M* is invertible if the parameters are positive. In order to obtain the *i*′’th component of *Ĝ*(*x̃*), consider the *i*’th entry of the canonical basis *b_i_*, compute (*L*^(0)^)^+^*b_i_*, and identify the component that is in *H***0**. E.g., for *i* = 1 we find *b*_1_ = *ṽ*^2^, and (*L*^(0)^)^+^*ṽ*^2^ = –2*ṽ* [(–*ζ*/*ϑ*)*ỹ* + *γϑκṽ*] + 2*ζκ*, where –2*ṽ* [(*–ζ*/*ϑ*)*ỹ* + *γϑκṽ*] ∈ *H*_2_ and 2*ζκ* ∈ *H*_0_. Hence, (*Ĝ*(*x̃*))_1_ = 2*ζκ*. The equation (*L*^(0)^)^+^(*M*^−1^*h*) = *h* + *g*(*x̃*) with *g*(*x̃*) = *Ĝ*(*x̃*) *M*^−1^*h* implies

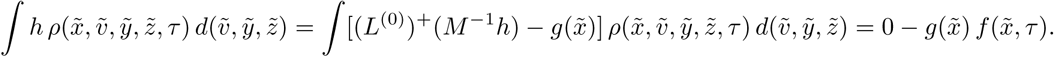

Solving the system for *h*(*x̃*, *ṽ*, *ỹ*, *z̃*) = *γ*/2*κ ṽz̃*, we obtain

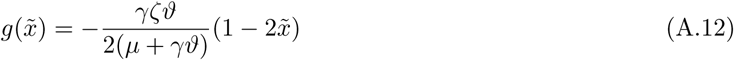

by using the computer algebra package MAXIMA (Maxima, 2014). In summary, we have

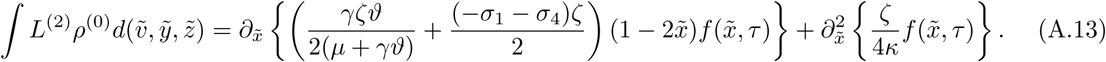

Now, we turn to the computation of the first integral in (A.9). With

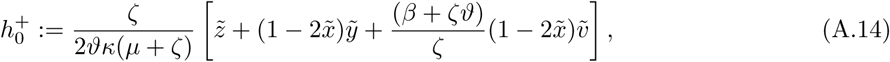

we obtain 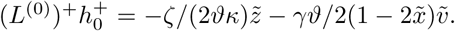 Remember that full derivatives w.r.t. *ṽ*, *ỹ* or *z̃* vanish upon integration. So,

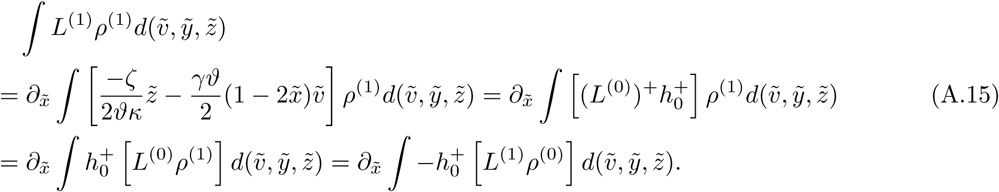

To handle 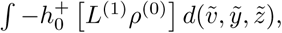 we use partial integration to move all derivatives in *L*^(1)^ w.r.t. *ṽ*, *ỹ*, *z̃* from 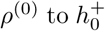 (note that 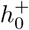 is linear in these variables, s.t. all second derivatives in these variables vanish). Let us consider one of these terms occurring in 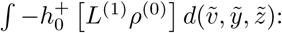

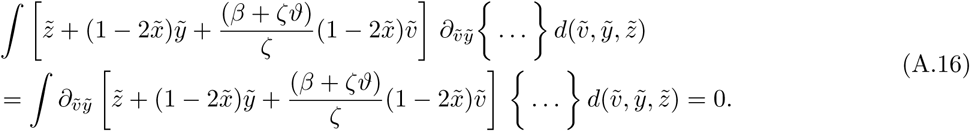

It is not possible to use the same procedure for *x̃*, as we do not integrate w.r.t. *x̃*. Here we use the product rule, e.g.,

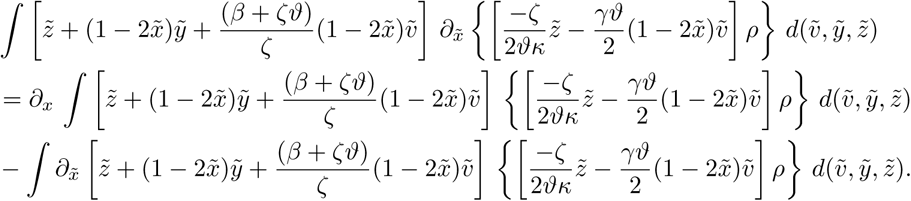

In this way, we obtain

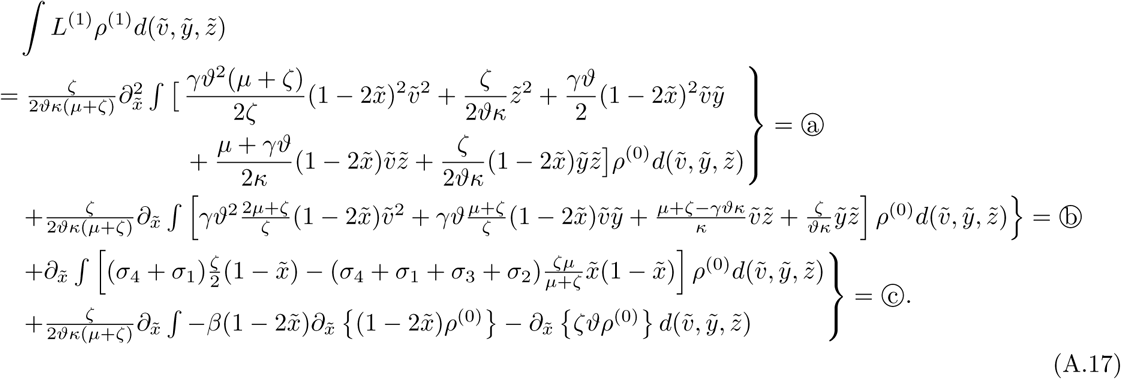

We will treat all three terms separately starting with Ⓒ.

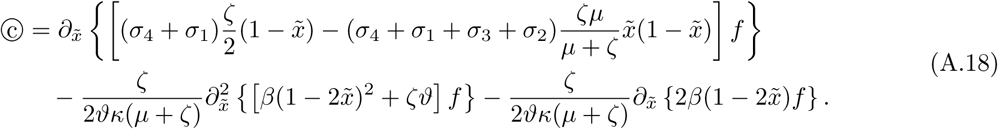

To compute ⓑ, we proceed as in the computations for ∫ *L*^(2)^*ρ*^(0)^*d*(*ṽ*, *ỹ*, *z̃*), that is, we solve

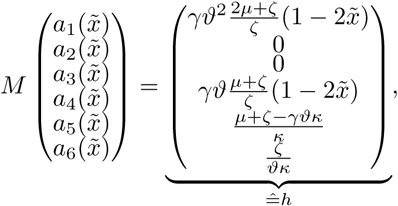

and find

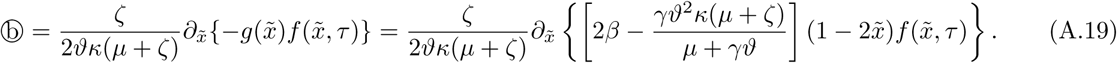

For ⓐ, this can be done similarly, where the system to be solved is

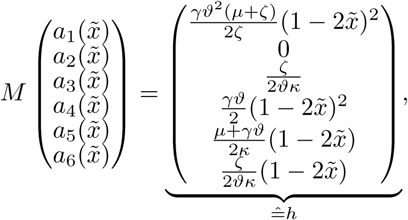

and as a result, we have

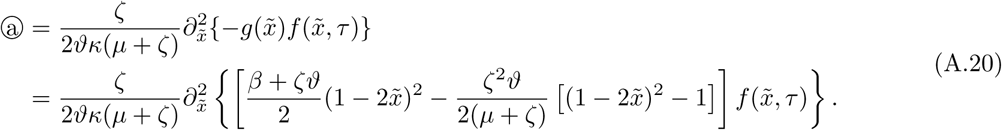

With ∫ *L*^(1)^*ρ*^(1)^*d*(*ṽ*, *ỹ*, *z̃*) = ⓐ + ⓑ + ⓒ and

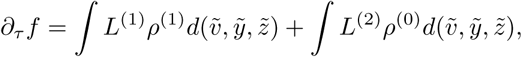

we obtain

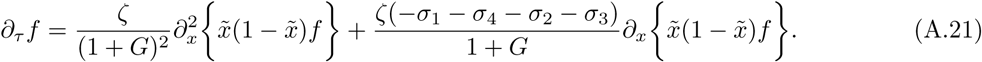

## A.3. Fixed population size and fluctuating seed bank

We only present the main steps for this and the following model, as the computations are lengthy and resemble that of the last model.

Let *p*_*i*,*j*,*k*_ (*t*) = ℙ(*X_t_* = *i*, *Y_t_* = *j*, *Z_t_* = *k*) be the probability distribution of the resulting stochastic process. The corresponding master equation reads:

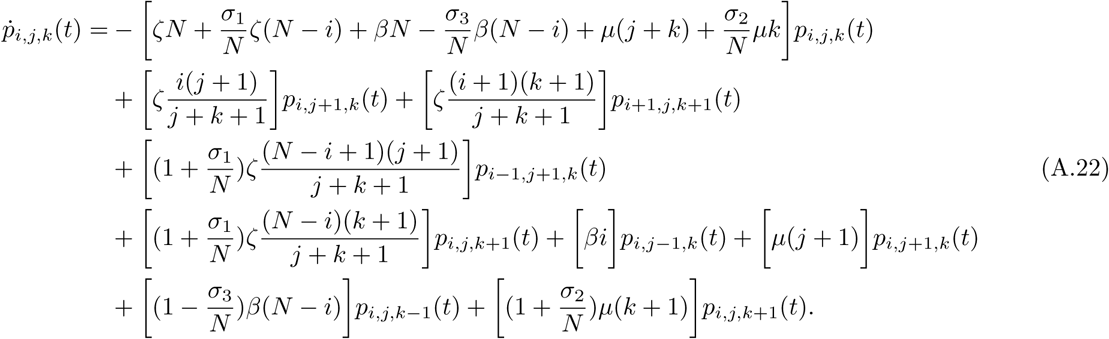

We now obtain the Fokker-Planck equation for the approximating diffusion process as described before. The first step is to transform the system to a quasi-continuous state space by scaling with *N*^−1^, *i.e.*, defining *x* = *i*/*N*, *y* = *j*/*N*, *z* = *k*/*N*, *h* = 1/*N* and the quasi-continuous density *u*(*x*, *y*, *z*, *t*) = *p*_*i*,*j*,*k*_(*t*). The resulting PDE reads

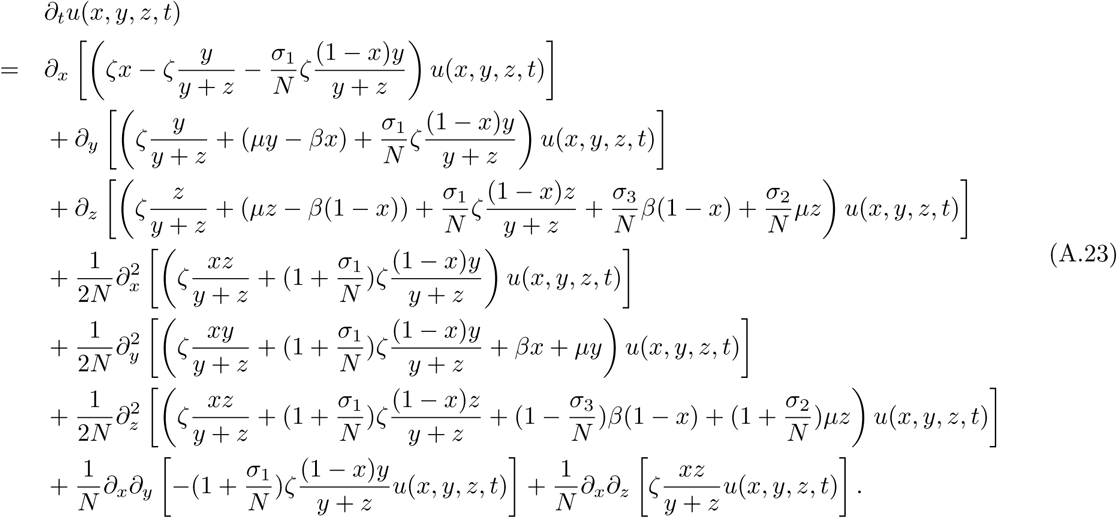

For *N* → ∞, we obtain a deterministic model, governed by the ODEs

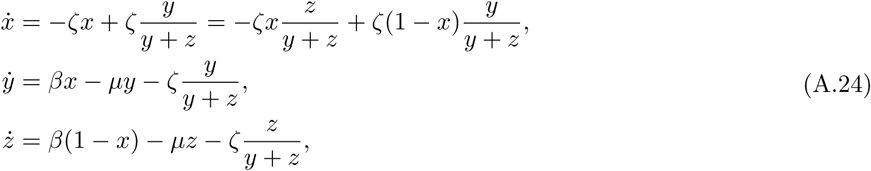

which is a neutral competition model since the selection terms vanish in the limit.

### Proposition A.3.

*Let ϑ* = (*β* – *ζ*)/*μ* > 0. *Then*, *there is a line of stationary points for (A.24) in* [0, 1] × 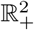 *given by*

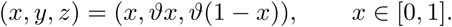

*The line of stationary points is transversally stable. The eigenvectors perpendicular to the line of stationary points (together with the eigenvalues) read*

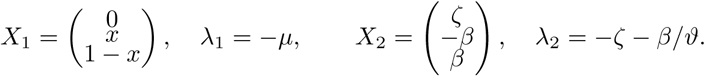

The proof of this proposition is straight forward, along the lines of the proof of proposition A.1. To formulate the inner solution, we introduce local coordinates

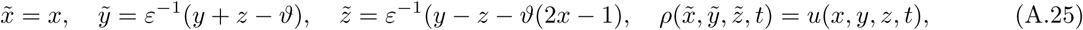

where *ϑ* = (*β* – *ζ*)/*μ* again and *ε*^2^ = 1/*N*. Alternatively formulated, we have

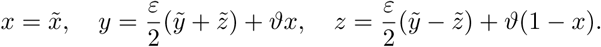

*ỹ* can be thought of as measuring the deviation of the total amount of seeds from its deterministic value *ϑ* and *z̃* as measuring the deviation of the allele ratio in seeds from the allele ratio in plants. Both are scaled by *ε*^−1^ so we expect them to be of order 𝒪(1). By transforming derivatives, we have

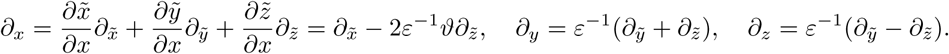

We now approximate *∂_tρ_*(*x̃*, *ỹ*, *z̃*, *t*) by transforming all terms on the r.h.s. of (A.23) and ignoring terms of 𝒪(*ε*^3^). We find

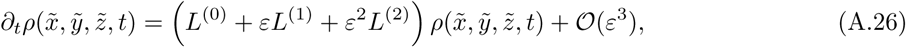

with linear differential operators *L*^(0)^, *L*^(1)^ and *L*^(2)^ that take the form

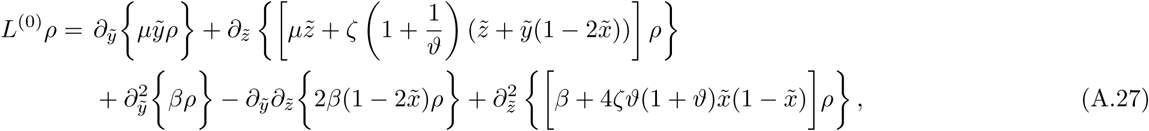

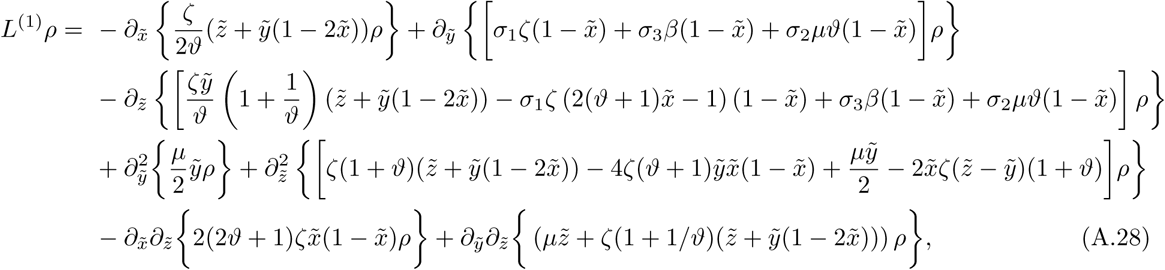

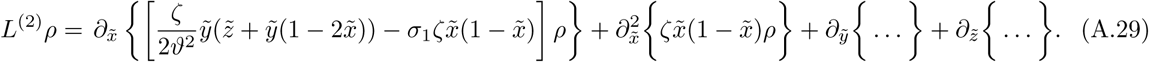

Before we proceed, we define the functions

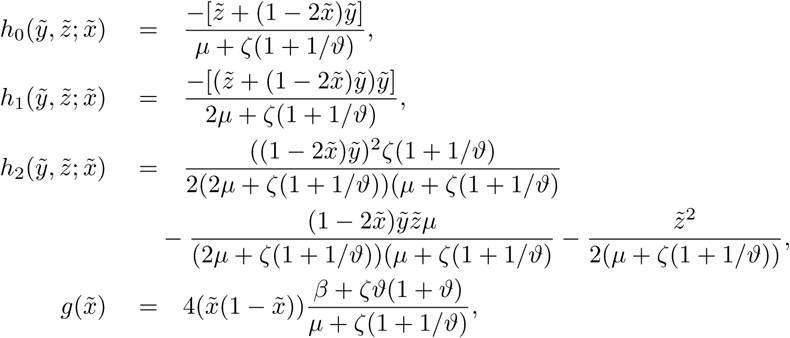

and note that

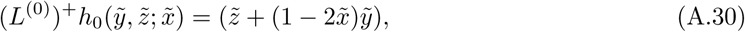

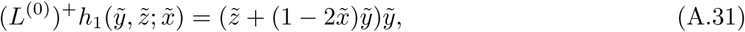

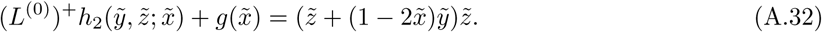

That is, *h*_0_ and *h*_1_ are eigenfunctions of (*L*^(0)^)^+^. Observing the system on slow time *τ* = *ε*^2^*t* = *t*/*N*, we see rapid dynamics for *τ* close to zero, *τ* ∈ 𝒪(1/*N*) to be precise. This is the new boundary layer. For *τ* ∈ 𝒪(1), only slow drift effects along the equilibrium line should remain. Being interested in the outer solution that develops on the slow time scale, we make the Ansatz

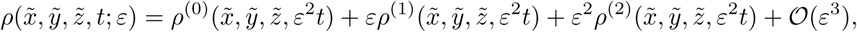

assuming *∂_tρ_* = 𝒪(*ε*^2^) to focus on the outer solution and neglect the boundary layer. We plug this Ansatz into (A.26), compare same order terms on both sides and obtain

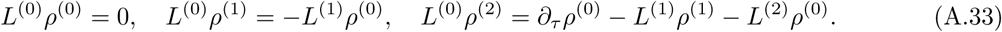

We have 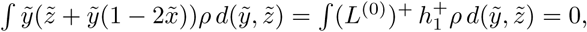 and hence

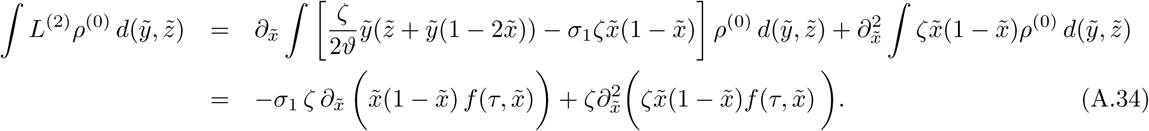

Using the same procedure as in Appendix A.2, consider eqn. (A.15)-(A.17), to handle ∫ *L*^1^ *ρ*^1^ *d*(*ṽ*, *ỹ*, *z̃*), and by applying (A.30) - (A.32), we find

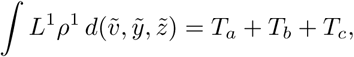

where

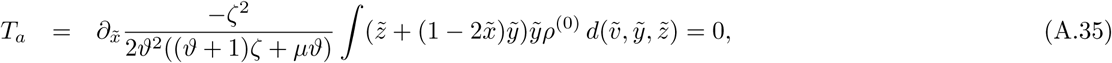

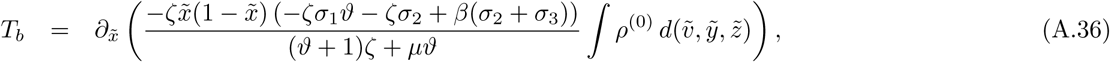

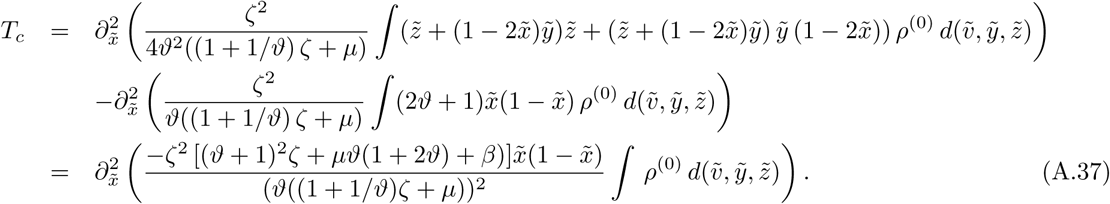

With

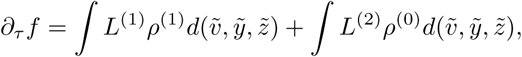

we find

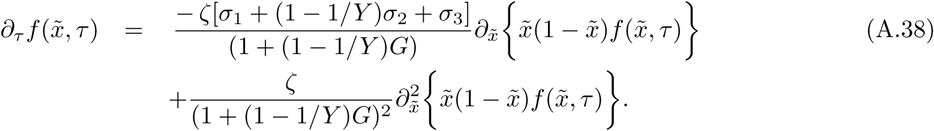

## A.4. Logistic population dynamics and fluctuating seed bank

Let *p*_*i*,*j*,*k*,*l*_(*t*) = *P*(*X*_1,*t*_ = *i*, *X*_2,*t*_ = *j*, *Y_t_* = *k*, *Z_t_* = *l*) be the probability distribution of the resulting stochastic process. The corresponding master equation reads:

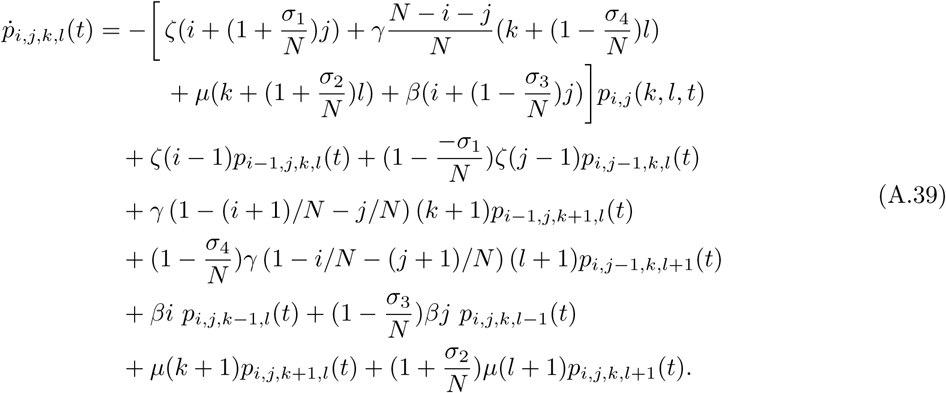

We transform the system to the quasi-continuous state space with rescaled parameters *x*_1_ = *i*/*N*, *x*_2_ = *j*/*N*, *y* = *k*/*N*, *z* = *l*/*N*, *h* = 1/*N* and quasi-continuous density *u*(*x*_1_, *x*_2_, *y*, *z*, *t*) = *p*_*i*,*j*,*k*,*l*_(*t*). After expanding about (*x*_1_, *x*_2_, *y*, *z*) in terms up to order 𝒪(*ε*_2_), we have a 4-dimensional Fokker-Planck equation again:

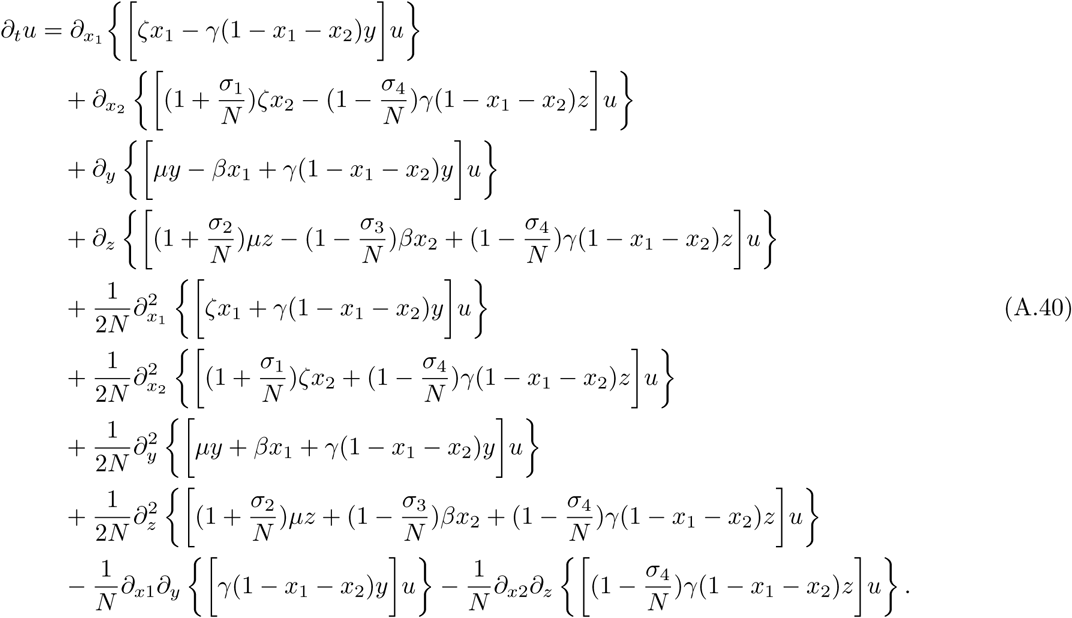

The limiting deterministic system for *N* → ∞ has dynamics

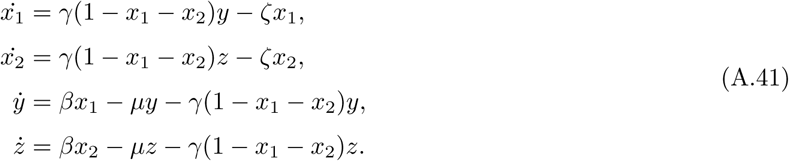

As before, we can show that a line of stable equilibria exists:

### Proposition A.4.

*Let ϑ*:= (*β* – *ζ*)/*μ* > 0, *κ*:= (*γϑ* – *ζ*)/*γϑ* ∈ [0, 1]. *Then*, *there is a line of stationary points in* [0, *κ*]^2^ × 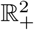 *given by*

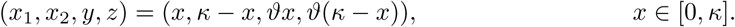

*The line of stationary points is transversally stable (locally and globally).*

Since the proof is similar to the proof of Proposition A.1, it is omitted here.

## A.4.1. Perturbation approximation

As before, new local variables for the boundary layer around the equilibrium line are defined:

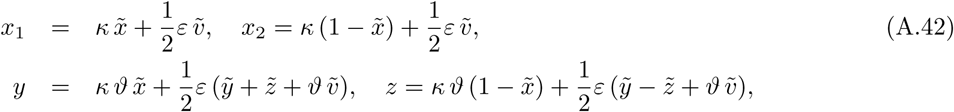

where

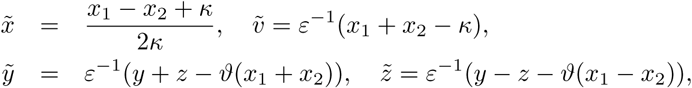

and *ε*^2^ = 1/*N*. The considerations for the global behavior are the same as for model 2. We choose the same local variables for the boundary layer around the equilibrium line and transform (A.40) up to terms of 𝒪(*ε*^3^) and higher. The resulting reparametrized Fokker-Planck equation in the local variables is

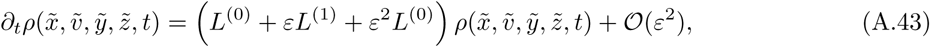

with linear differential operators

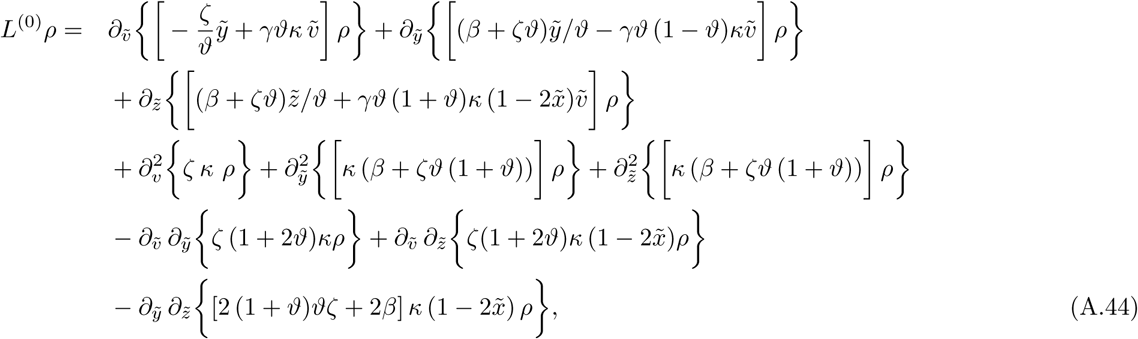

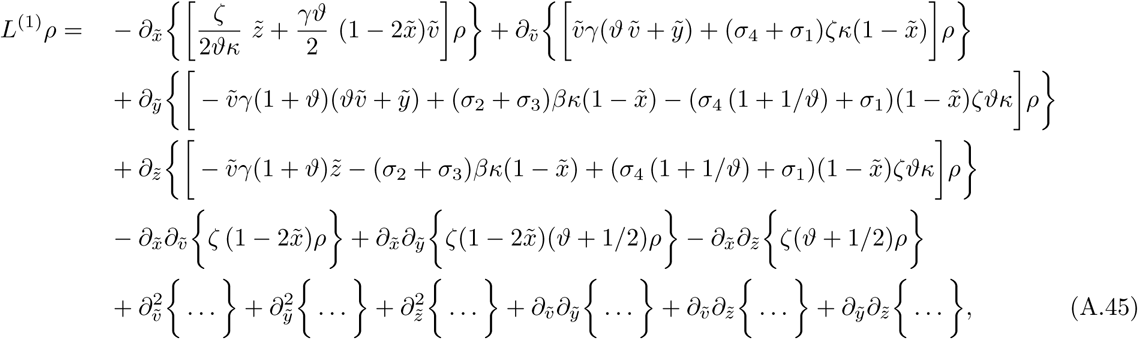

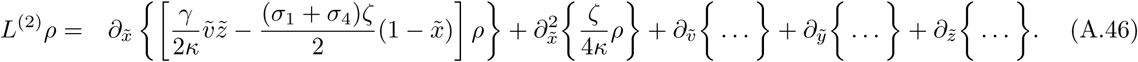

To handle the integrals below, we define an operator *M*: *H*_2_ → *H*_0_ resp. *Ĝ*: *H*_2_ → *H*_0_ in a similar way as above (Proposition A.2). Recall that we use (*ṽ*^2^, *ỹ*^2^, *z̃*^2^, *ṽỹ*, *ṽz̃*, *ỹz̃*) as the basis in *H*_2_. The operator *M* has the representation

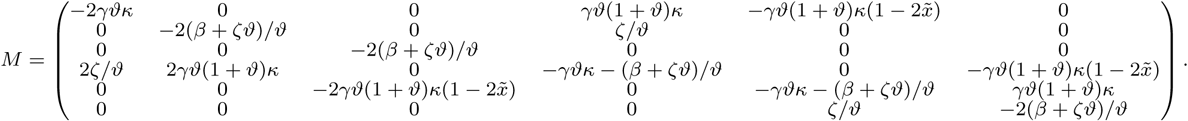

If we define *g* via *g*(*x̃*) = *Ĝ*(*x̃*) *h*, we find

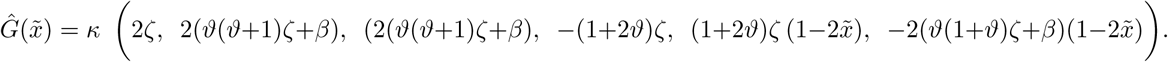

As before, we employ time scale separation and focus on a solution evolving on the slow time *τ* = *ε*^2^*t* = *t*/*N*, using the Ansatz

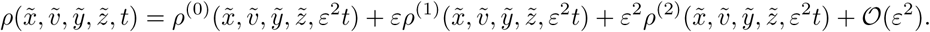

If we define 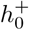 by

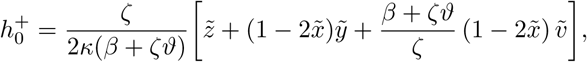

we find

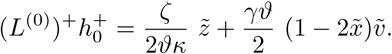

By now, we have all ingredients together to go along the same route as in Appendix A.2. We start with ∫ *L*^(2)^*ρ*^(0)^ *d*(*ṽ*, *ỹ*, *z̃*). If we use

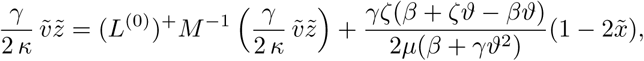

we obtain

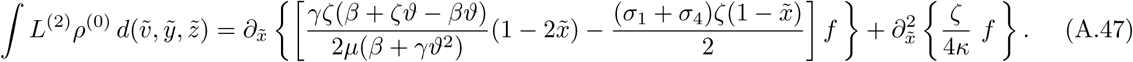

Next, we turn to ∫ *L*^(1)^*ρ*^(1)^ *d*(*ṽ*, *ỹ*, *z̃*) = 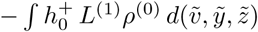 with

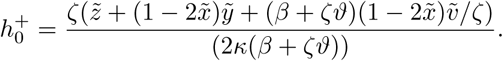

We integrate by parts w.r.t. all derivatives *∂_ṽ_*, *∂_ỹ_*, *∂_z̃_*, and respectively move the derivatives *∂_x̃_* in front of the integral by means of the chain rule, so that we obtain

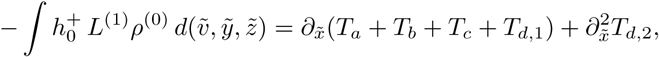

with

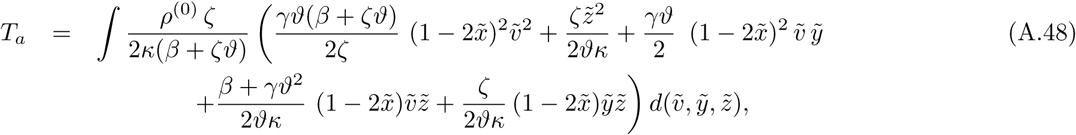

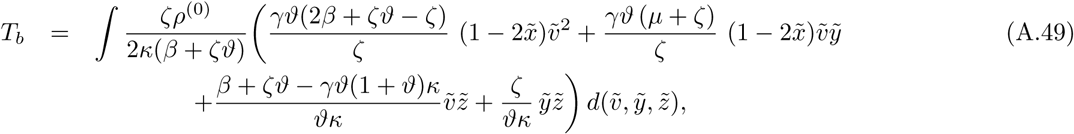

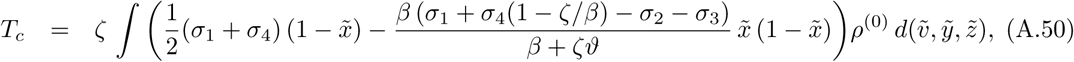

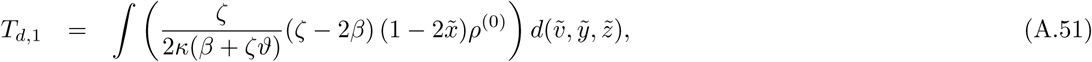

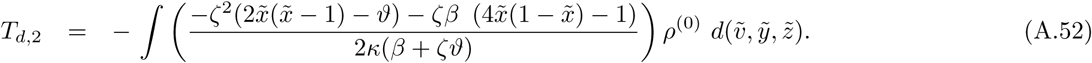

In particular, the integrals in *T_a_* and *T_b_* can be transformed using the operators *M* and *Ĝ*,

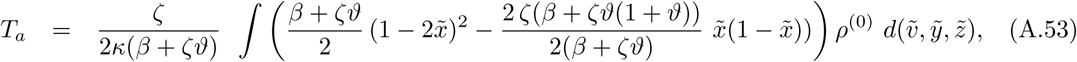

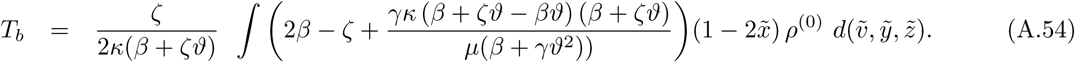

Recall that in the present variant of the model *ϑ* = (*β* – *ζ*)/*μ*, *G* = *ζ*/*μ* and *Y* = *β*/*ζ*. With

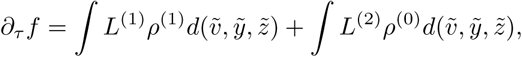

we find

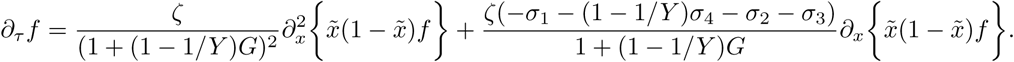

